# Structural Aspects of Homologous DNA-DNA Interactions Revealed by Partitioning of RIP Mutations

**DOI:** 10.1101/344036

**Authors:** Alexey K. Mazur, Eugene Gladyshev

## Abstract

In some fungi, a process known as Repeat-Induced Point mutation (RIP) can accurately identify and mutate nearly all gene-sized DNA repeats present in the haploid premeiotic nuclei. Studies of RIP in *Neurospora crassa* have suggested that the sequence homology is detected between intact double helices without strand separation and participation of RecA homologs. These studies relied on the aggregated number of mutations as a simple quantitative readout of RIP activity and did not try interpret the distributions of mutations along DNA. Important additional information can be extracted by transforming these distributions into profiles of a new parameter called partitioned RIP propensity (PRP) which takes into account the site density as well as the sequence context. This approach revealed surprising systematic variations of PRP due to the position of a given DNA segment relative to the homology boundaries and its topology. Notably, identical pairs of direct versus inverted repeats produce very distinct PRP profiles. This effect could be rationalized assuming a specific redistribution of the supercoiling stress produced by the previously discovered untwisting of paired of DNA homologs. Similar mechanisms account for other persistent features of PRP profiles, and this general topological model raises an intriguing possibility that local DNA supercoiling provoked by homologous dsDNA-dsDNA pairing can modulate the overall structure and properties of repetitive DNA. These effects can be particularly strong in the context of long tandem repeat arrays that are typically present at the (peri)centromeric regions of chromosomes.

## INTRODUCTION

Homologous recombination-independent pairing of chromosomes (or individual chromosomal loci) represents a prominent feature of eukaryotic biology. Since its first report in *Diptera* insects more than a century ago [1], such pairing was documented in many biological systems, including partial or complete pairing of homologous chromosomes during the first meiotic prophase in mice [2], round worms [3], fruit flies [4], and fungi [5] [6] as well as transient pairing of homologous loci in somatic nuclei of mammals [7, 8] and budding yeast [6]. These microscopy-based observations were recently extended by high-throughput genomic studies in budding yeast [9] and animals [10], which reported a subtle but general propensity of homologous sequences to colocalize in the nucleus. The proposed models of recombination-independent pairing invoke either indirect (protein-mediated, e.g., [11], RNA-mediated, e.g., [12], or transcription-mediated [13]) or direct (physical pairing of intact, double-stranded DNA molecules, e.g., [14]) interactions. Yet, despite the large number of such models, the mechanistic nature of a general process that could promote recombination-independent pairing has remained elusive [15].

The repertoire of homology-dependent phenomena also includes a poorly understood process of somatic, RNAi-independent Repeat-Induced Gene Silencing (RIGS). During RIGS, newly integrated repetitive DNA sequences can trigger the formation of heterochromatin in organisms as diverse as plants [16], mammals [17] and filamentous fungi [18, 19]. While in some cases heterochromatic silencing can be guided by dedicated RNA interference (RNAi) pathways [20, 21], in both mammals and filamentous fungi RIGS is also known to occur in the absence of RNAi [22, 23], suggesting that repetitive DNA may be recognized for silencing by a conserved yet unknown mechanism.

The best evidence for the existence of direct pairing interactions between intact homologous double-stranded DNA molecules is provided by the phenomenon of Repeat-Induced Point mutation (RIP), which was first described in the filamentous fungus *Neurospora crassa* [24], and which is now known to occur in many fungi, specifically in the haploid premeiotic nuclei that continue to divide by mitosis in preparation for karyogamy and subsequent meiosis [25]. During RIP, segments of genomic DNA that share more than a few hundred base-pairs of sequence homology are detected and mutated by numerous cytosine-to-thymine (C-to-T) transitions [24–26]. Importantly, these segments can be targeted by RIP irrespectively of their specific sequence composition, coding capacity, or positions (absolute as well as relative) in the genome. Recent work has shown that such general and highly efficient repeat-sensing process occurs normally in the absence of Rad51 (or Dmc1), which mediate recognition of DNA homology for recombinational DNA repair in eukaryotes [26, 27]. The critical insight into the mechanism of RIP was based on the finding that the number of induced C-to-T transitions could, in principle, accurately reflect the amount of inducing DNA homology [26]. Importantly, the unexpectedly accurate relationship between the amount of homology and the number of ensuing mutations was observed in a genetic system that permitted systematic manipulation of the assayed repeats. This clarified a number of basic questions that would normally be studied only *in vitro* [27]. Specifically, it was found that RIP could detect the presence of homologous trinucleotides interspersed with a matching periodicity of 11 or 12 base-pairs along the participating chromosomal segments. Attempts to disrupt this apparently rigid periodicity abrogated RIP [26]. Another key finding was that the sequence composition of homologous triplets strongly affected the repeat recognition for RIP [28]. These and other results suggested that the homology sensing in RIP involved co-aligned double-stranded DNA helices that interacted using individual triplets of homologous base-pairs no more than one triplet per each helical turn [26, 27]. The steric possibility of direct homologous dsDNA pairing consistent with the above constraints was found in silico by using methods of quantum chemistry and molecular dynamics [29]. In this model, two aligned dsDNA pair by major groove surfaces to form short quadruplexes between homologous (self-complementary) base-pair triplets once per a few helical turns. The same type of pairing was considered in the early theories of DNA replication [30] and homologous recombination [14]. The computations predicted a significant difference in the stabilization energies of AT and GC quartets, with a much stronger pairing for quadruplexes formed between GC-rich DNA.

RIP has long been known to require a specialized C5-cytosine DNA methyltransferase (DMT) RID (RIP defective) [31]. This DMT has a compact structure that includes a signature of the BAH-like domain, a module implicated in protein-protein interactions in the context of silenced chromatin [26]. While RID-like DMTs represent an ancient protein family [32], they only occur in filamentous fungi, suggesting that RIP itself might be a phylogenetically restricted phenomenon. Intriguingly and contrary to this idea, recent results suggest that RIP may be a manifestation of the general phenomenon of heterochromatic silencing of repetitive DNA [33, 34]. Such dsDNA-dsDNA interactions may be important in regulating the epigenetic state of tandem repeats even in organisms that do not feature the canonical heterochromatic H3K9me2/3 and H3K27me2/3 [35, 36]. These results specifically implicate the SUV39 lysine methyl-transferase DIM-5 (which catalyzes all trimethylation of histone H3 lysine 9 residues in *N. crassa*) as a mediator of RID-independent RIP, which also requires DIM-2 (the second DMT in *N. crassa*) [34, 37]. Importantly, both RID-dependent and RID-independent pathways of RIP are triggered by the same homology signal in *N. crassa*, but, to the first approximation, appear independent from one another [33]. Taken together, the previous studies have demonstrated that the phenomenon of RIP can serve as a powerful quantitative model of homology-dependent epigenetic processes, which can shed light on both the eukaryotic phenomenon of repeat silencing and the nature of physical mechanisms of homologous dsDNA-dsDNA interactions.

The previous studies used the bulk number of RIP mutations as a sole quantitative measure of RIP activity, and did not take into account the additional information involved in the distribution of mutations along the assayed repeat constructs. Here we present a computational approach for extracting and evaluating such information. This method takes raw persite counts of mutations and computes a partitioned RIP propensity (PRP) which evaluates the exposure to RIP for a cytosine site in the vicinity of any position along DNA. It characterizes a small DNA segment rather than a particular site or sequence motif and more accurately distinguishes regions that are not equivalent with respect to RIP mutation. The PRP profiles of new synthetic DNA constructs as well as those computed from previous data revealed surprising general features of RIP that had been previously obscured by the inherently noisy nature of point mutations. Several of them, notably, a striking qualitative divergence between the PRP profiles for identical pairs of closely-positioned repeats in either direct or inverted orientation, were attributed to distinct patterns of supercoiling generated due to dsDNA-dsDNA pairing and small untwisting of the paired duplexes. We also confirmed that, in qualitative agreement with theoretical predictions [29], AT-rich repeats trigger RIP less efficiently than those with normal AT content. Collectively, our results advance the idea that RIP (and by extension RIGS) involves direct pairing of double-stranded DNA molecules by the formation of interspersed quadruplexes. Our findings may have important implications for understanding the mechanisms of other pairing phenomena that involve recombination-independent recognition of DNA homology. These results may also be relevant to understanding the three-dimensional organization and function of long tandem repeats that are typically found in (peri-)centromeric regions of mammalian chromosomes.

## MATERIAL AND METHODS

### Experimental data

The experimental data analyzed below were all taken from the previous reports [27, 33], with one additional test carried out by using the same protocol. The corresponding procedures including the plasmid construction and the primer sequences are described in the Appendix.

### Software availability

The algorithm for building PRP profiles is implemented in C. The source code will be freely available from GitHub.

## RESULTS

### Partitioned RIP propensity profiles

RIP mutation represents an intrinsically stochastic process, yet its general tendency can be revealed by analyzing a set of individual products sampled from a large population of the cross progeny. A representative example of the experimental RIP dataset (based on 24 randomly sampled RIP products [27]) is shown in Fig. 1A. The longitudinal variation in mutation frequency carries information that, in principle, can be used to gain further insights into the mechanism of DNA homology recognition for RIP. However, the mutation frequency of a given DNA segment also depends on the number of internal cytosine sites and their sequence context, which overshadows other contributing factors and makes this value unsuitable for further analysis. To circumvent these problems, we introduce a partitioned RIP propensity (PRP) defined as the probability of RIP for cytosine sites of a particular type in a given short segment of genome DNA. The persite probability of RIP is estimated as *P* = *M*/*N* where *M* is the number of mutations per cross and N is the number of sites. Evidently, *P <* 1 and *P* → 1 with exhaustive RIP. The sequence context is taken into account in the nearest-neighbor approximation. A strong dependence upon the identity of the cytosine neighbors in the DNA stack is expected for the methylation step because the involved DMTs have to pull the targeted nucleotide out from the double helix [32]. Indeed, in *N. crassa*, RIP preferentially affects cytosines in CpA dinucleotides [38]. Fig. 1B indicates that the quantities of cytosine sites with different next neighbors strongly and asynchronously fluctuate along DNA, and that these fluctuations cannot be neglected. At the same time, the effect of the next nearest neighbors is much less significant. Examination of our earlier data [27] confirms that for RIP of CpA steps the identity of the previous and next nearest neighbors have minor effects (see Appendix). Therefore we partition all cytosines into CpA, CpT, CpG, and CpC dinucleotides and assume that in the first approximation the intrinsic mutation rates do not depend upon other neighbors. The corresponding probabilities are denoted as *P*_CN_ where N stands for the second base.

**FIG. 1:**
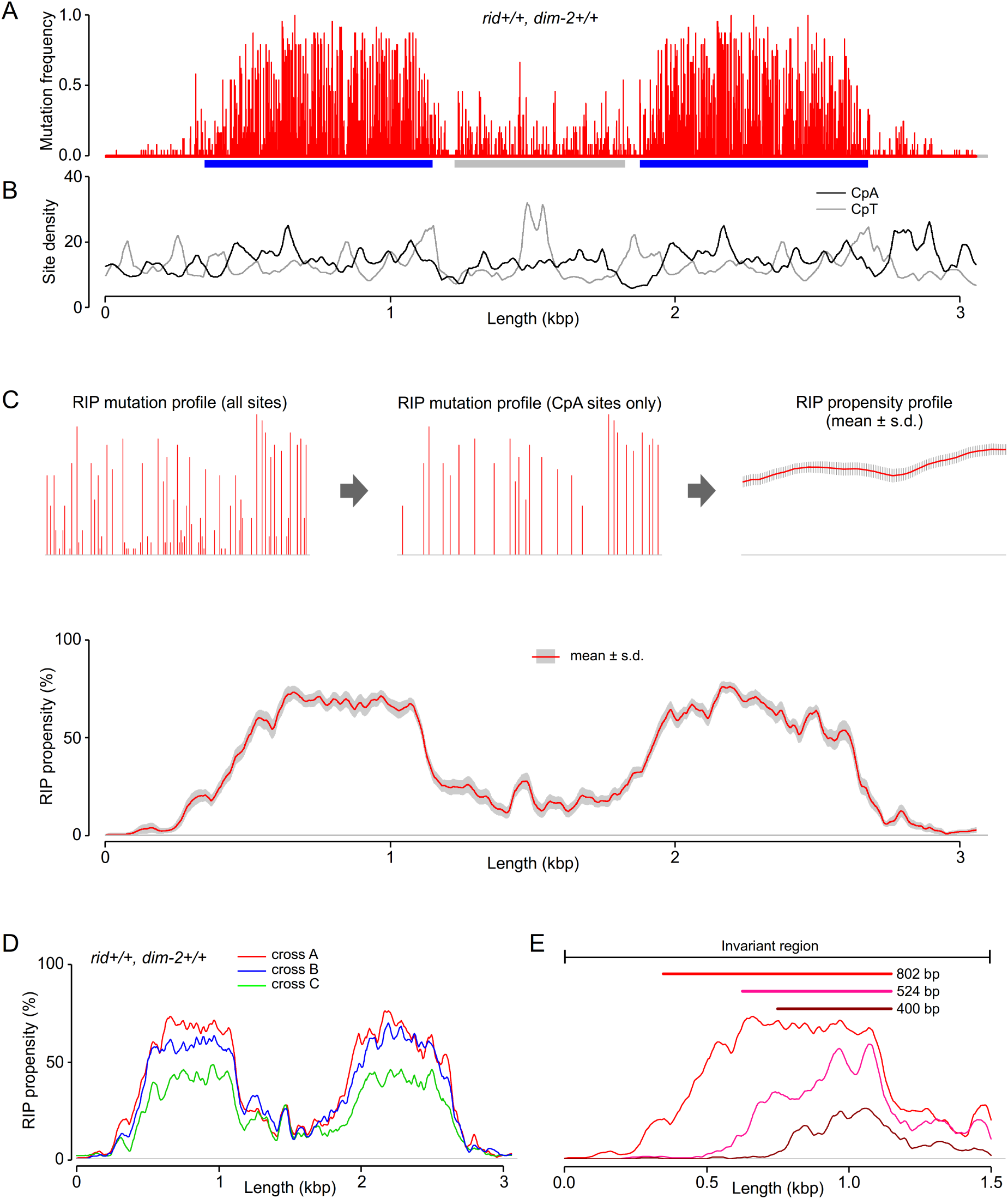
Construction of partitioned PRP profiles. (A) Experimental mutation frequencies for a 802 bp direct repeat construct from the previous study [27]. The repeats and the linker are marked by the blue and gray bars, respectively. (B) Computed site densities for two cytosine base-pair steps. (C) Construction of the CpA PRP profiles from the data in panel A. (D) Three PRP profiles obtained in independent genetic experiments for the same 802 bp repeat construct. (E) Three PRP profiles for the same DNA, with different segments repeated in the right flanking region beyond 1.5 kbp. Computed from the data from the previous study [27]. The repeated segments are marked on top.

Any position s on the DNA molecule can be characterized by the density of cytosine sites *D*(*s*), the corresponding density of mutations *M*(*s*), and the probability density of RIP

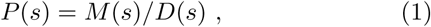

which gives a continuous profile on a long DNA. *P*(*s*) evaluates the exposure or the propensity to RIP for a cytosine site in the vicinity of position *s*, that is, it characterizes a small DNA segment rather than a particular site or a sequence motif. As noted above, the site densities fluctuate asynchronously and with similar amplitudes (Fig. 1B). The strong sequence dependence of RIP does not allow mixing of different types of cytosine sites in Equation (1). Instead one has to use a partitioned propensity (PRP) computed for a particular type of cytosine sites. In the further discussions we use only the PRP profiles for CpA steps obtained as *P*_CA_(*s*) = *M*_CA_(*s*)/*D*_CA_(*s*), therefore, the subscript will be omitted. Additional details concerning the construction of PRP profiles can be found in the Appendix.

Fig. 1C displays the computational steps and the result obtained from the data in Fig. 1A. The bottom panel shows the average profile with standard deviations obtained by bootstrap resampling with replacement. The profile shape reflects that of the bar plot of mutation frequency in Fig. 1A, and also reveals the features of RIP mutation that have previously been obscured by the local sequence effects. The estimated error (Fig. 1C) is relatively large, suggesting that some of the fine profile features may be irreproducible. Contrary to this expectation, we find that even such small-scale features are very robust. Specifically, we find that these features can be recapitulated by the two additional, previously published experiments [33] involving the same repeat construct. As analyzed in Fig. 1D, all small-scale peaks, which appear to be within the experimental error in Fig. 1C, are consistently revealed by the two independent profiles (each based on 24 random RIP products).

The evident contradiction between the apparent error and the robustness of fine details can be explained as follows. Suppose we have a set of identical normalized profiles scaled with a random factor. Despite identical profile shapes, bootstrapping of such data would give a non-zero estimate of errors which, however, refers to the fluctuations of the scaling factor rather than the shape of the profile. This feature is an intrinsic property of the experimental RIP data obtained by sequencing randomly sampled RIP products. The total number of mutations detected in a given DNA molecule is determined, among other factors, by the number of RIP cycles that this particular molecule has passed through. This value strongly varies in any RIP experiment. In the DNA samples used in Fig. 1A, the integral RIP intensity varied by more than a factor of two. Whatever the biological mechanisms responsible for these variations, they do not affect the fine structures of PRP profiles, which explains their robustness in repetitive experiments.

One more representative example is shown in Fig. 1E. These three profiles were obtained for the invariant part of DNA in the series of constructs that were used earlier to study the length dependence of RIP in *N. crassa* [27]. The upper profile is identical to the left part of the red profile in Fig. 1D. The variable right part with the second repeat copy is not shown. The repeats were gradually shortened as indicated in the figure. In the 400 bp interval shared by all these repeats the profiles exhibit two prominent peaks at similar positions. These features on PRP profiles should be interpreted cautiously. For instance, peaks and valleys do not necessarily correspond to the accessibility of DNA to the involved DMTs because the corresponding profiles may result from superposition of a large number of dsDNA-dsDNA pairing modes with different axial rotations. Nevertheless, they provide valuable information by showing that different DNA segments along the repeat construct are not equivalent with respect to RIP mutation, and the above peaks in Fig. 1E can be due to specific structures formed during RIP at repeat ends. This figure also illustrates the RIP saturation effect (see also the Appendix). With the repeat length increased the RIP propensities approach 100%. As a result, the sequence-dependent peaks are smoothed and the profile takes the shape of a plateau.

Taken together, our analysis indicates that the PRP profiles have the potential to reveal the conserved, previously hidden features of the RIP process.

### PRP profiles in homologous pairs are not identical

In *N. crassa*, RIP mutation can be mediated by two molecular pathways that are triggered by the presence of DNA homology (Fig. 2A). The first, canonical pathway involves a specific DMT called RID (”RIPA defective”). Its analogs in other filamentous fungi are implicated in the vegetative and sexual development [39–41], but in *N. crassa* RID has no other known function [31]. The canonical RIP pathway predominantly results in mutations within the repeated sequences *per se*, rather than their flanking regions. The second, recently discovered pathway [33] involves DIM-2 (”Defective in methylation- 2”), which is normally responsible for all 5-cytosine methylation in *N. crassa* [42]. In addition to DIM-2 this pathway depends on several proteins known to be involved in the formation of heterochromatin (Fig. 2A). In the contexts of DNA constructs shown in Fig. 1 the DIM-2 pathway mainly results in mutations in the single-copy linker region between the repeats [33].

**FIG. 2:**
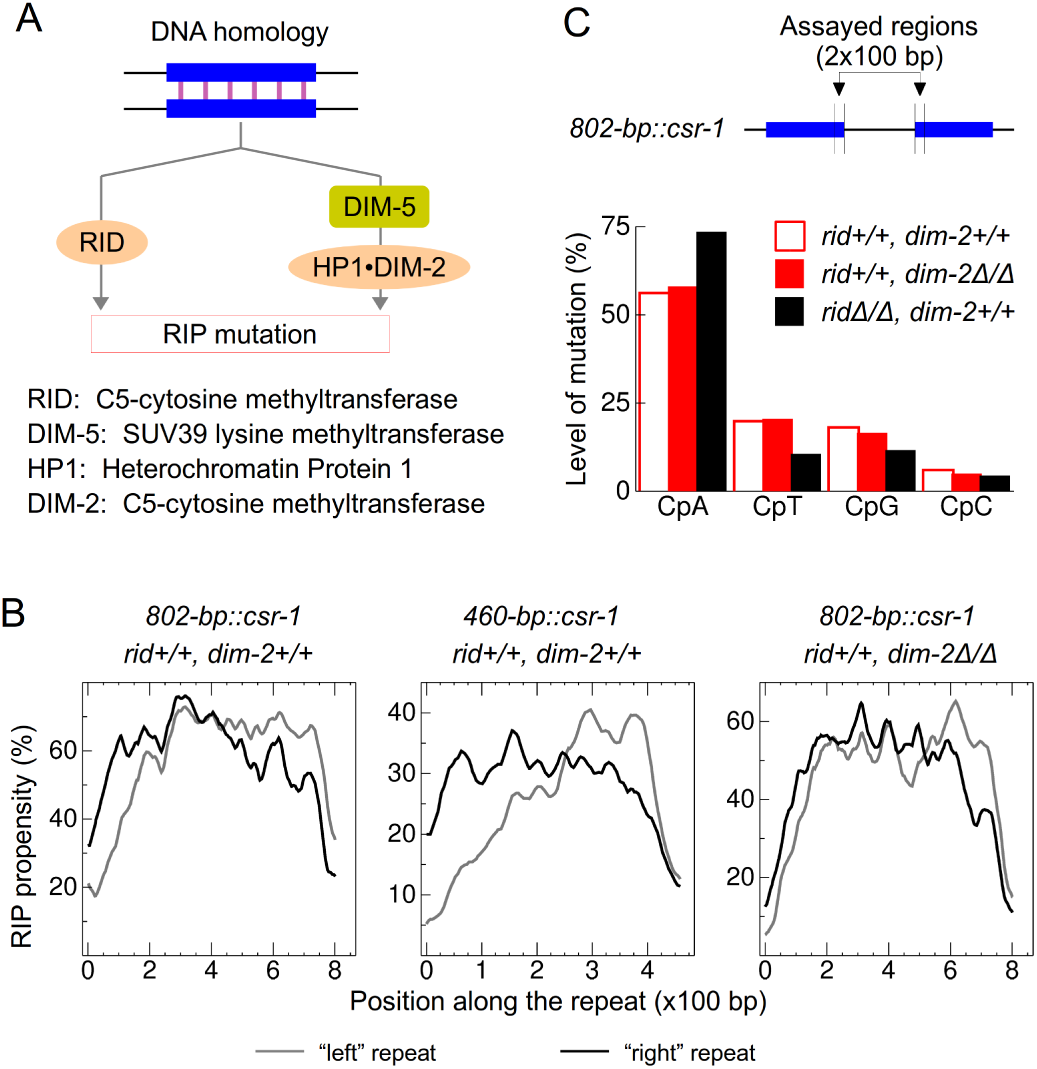
(A) Currently accepted model of the RIP process in *N. crassa*. Critically important protein components involved in the two pathways are indicated. (B) Aligned PRP profiles for three pairs of direct repeats with different lengths and genetic locations. The linker length was 706 bp. (A) The cytosine duplet composition of mutations effectively added in the 100 bp zones of the 802 bp repeats flanking the linker in the wild-type *N. crassa* (open bars) compared with the overall duplet compositions of RIP in the (rid+/+,dim-2Δ/Δ) and (ridΔ/Δ, dim-2+/+) strains.

It is understood that PRP profiles of identical sequences corresponding to pairs of recognized homologs do not have to match exactly because of the stochastic nature of point mutations. It turned out, however, that such profiles are always qualitatively distinct depending on their relative positions and orientations. Fig. 2B shows three representative examples for DNA constructs studied earlier [27, 33], with direct repeats separated by a linker of 706 bp. The left panel presents a pairwise comparison of PRP profiles corresponding to homologous DNA segments in Fig. 1A. The gray and black plots show the profiles for the left and right repeat copies, respectively. These are direct repeats, that is, the middle linker is on the right from the gray profile, and on the left from the black one. It is seen that the RIP propensity is higher in the flanking regions around the linker. At the same time, the aligned profiles have very similar fine structures, with many matching peak positions. In the middle panel, similar data are shown for another construct from the same series [27], with the repeat length reduced to 460 bp. Here the divergence between the gray and black curves looks more significant because of lower PRP levels. In fact, absolute deviations are similar and in both cases they involve regions of about 150 base-pair steps adjacent to the linker, which in the second case represents more than a half of the repeat length.

Since the DIM-2 dependent RIP on the linker spreads also into the repeats [33] one can reasonably suggest that this residual activity results in the increased RIP propensity in the DNA segments flanking the linker. Indeed, the divergence between the black and gray CpA profiles in the left panel of Fig. 2B is of the same order of magnitude as the RIP propensity estimated on the linker in Fig. 1C. This interpretation was checked by probing the same effect in the mutant (rid+/+,dim-2Δ/Δ) strain of *N. crassa*. As shown in the right panel of Fig. 2B, in the absence of DIM-2 the difference is still observed, therefore, the effect is definitely due to the main RID-dependent pathway. This conclusion is additionally supported by the results shown in Fig. 2C. It was noticed earlier that the sequence dependences of RIP in these two alternative pathways are somewhat different, with a stronger preference to CpA steps in the DIM-2-dependent RIP [33]. Fig. 2C examines the sequence composition of mutations that contributed in Fig. 2B to the additional RIP propensity in the regions flanking the linker in the 802 bp construct. It is seen that they resemble the characteristic pattern of RID rather than DIM-2-dependent RIP.

We conclude that the closeness of the linker somehow affects the RID-dependent RIP activity on the repeats, which increases the RIP propensity in the flanking regions of about 150 bp.

### Linker PRP profiles change with its length and repeat orientations

Fig. 3 compares PRP profiles obtained for three constructs in which a unique 337 bp sequence was repeated in different contexts. The data were taken from earlier experiments with the wild-type strain of *N. crassa* [27]. In the first construct (green) the repeats are in the direct orientation and separated by a linker of 706 bp. In the second construct (red) this linker was cut to 303 bp by removing its right-hand half. Comparison of these two profiles shown in the left and middle panels, respectively, reveals that, when the linker between direct repeats was reduced from 706 to 303 bp, the overall RIP intensity significantly increased. Interestingly, this increase in the middle of the linker is nearly two times larger than that on the repeats. These results might have been expected based upon other observations because (i) the probability to escape from RIP is larger when the two homologous sequences are widely separated in the genome, and (ii) the residual relative RIP activity beyond widely spaced repeats is negligible.

**FIG. 3:**
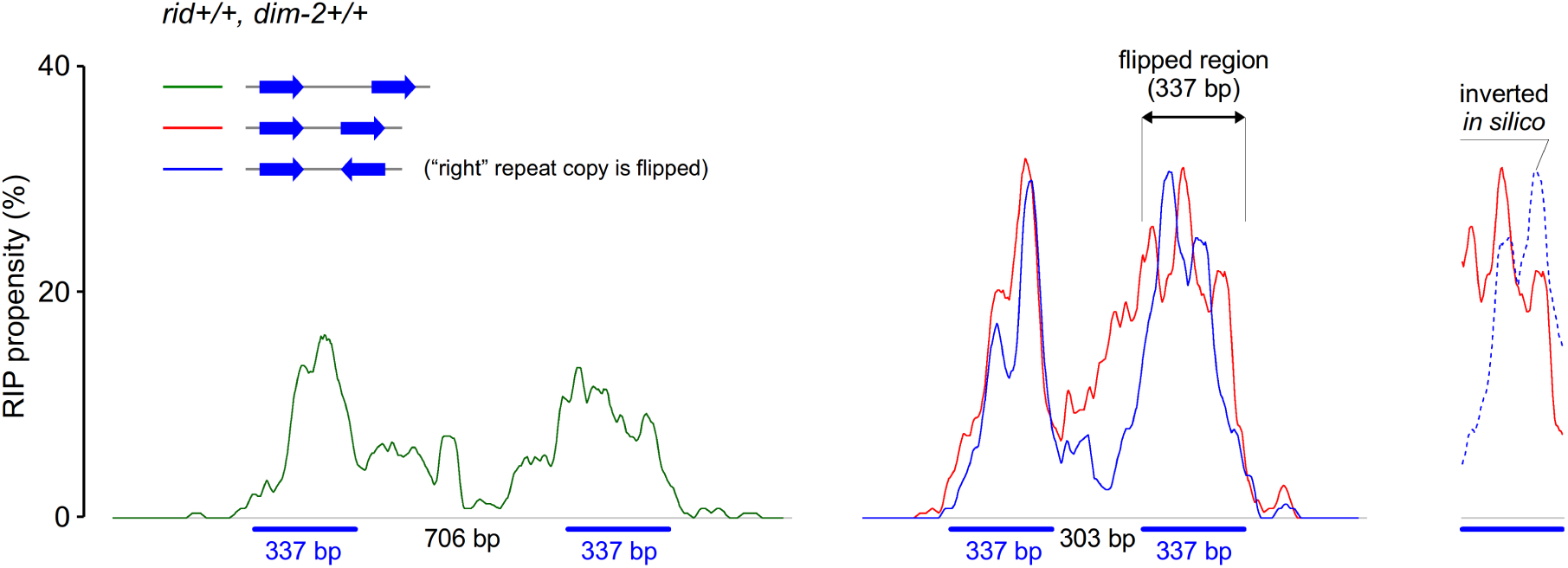
PRP profiles for the same repeated sequence in three different contexts. The repeat positions and orientations are shown by blue arrows and bars.

The third construct (blue) differs from the second one only by the inverted orientation of the right repeat copy. The PRP profiles for these two constructs are superimposed in the middle panel. Many features in these plots are evidently determined by the base-pair sequence. Notably, the invariant left repeat copy always produces similar profiles with the same positions of all peaks. With the right repeat copy inverted, the corresponding part of the PRP profile is also inverted. This is illustrated in the right panel of Fig. 3. It shows the segment marked by arrows in the middle panel, but with the blue trace flipped, which results in matching positions of all peaks because the base-pair sequences become aligned. For the linker, however, the two PRP profiles are strongly different, and the 303 pb DNA segment marked in the middle panel, which is invariant in the linkers of these three constructs, demonstrates very different properties even though its left flaking sequence is also invariant. In fact, the red and blue profiles of this segment in the middle panel differ so strongly that it is difficult to believe that they correspond to the same sequence, which suggests that they might be due to different RIP pathways.

Between the direct repeats (red) the RIP intensity is essentially similar to that on repeats, which is unusually high. Since the internal part of the linker is processed exclusively via the DIM-2-dependent RIP [33] this pathway should be specifically stimulated by some feature in this particular arrangement of homologous sequence segments. In contrast, between the inverted repeats the RIP intensity is lower than even in the mother (green) construct with the long linker. Moreover, this PRP profile exhibits a minimum in the middle of the linker, which was never found in the earlier studied constructs with direct repeats, but usual for the inverted orientation [27, 28]. A similar minimum in the middle of the linker is observed for RIP on direct repeats in the mutant (rid+/+,dim-2Δ/Δ) strain [33]. The blue PRP profile in the middle panel of Fig. 3A might be entirely due to the RID-dependent RIP because this activity can spread and include the boundary zones of short linkers. A quantitative comparison with the earlier data [33] indicates that in the (rid+/+,dim-2Δ/Δ) mutant this residual propensity is not very different from that inside the linker according to the blue profile in Fig. 3A.

The foregoing observations suggest that the DIM-2-dependent RIP on the linker is specifically stimulated by the direct repeat orientation and inhibited by the inverted one. We attribute these effects to the redistribution of the supercoiling stress inside the DNA loop formed by paired homologous segments. The possible mechanisms will be discussed in detail further below.

### Peaked structures of linker PRP profiles are not sequence-specific

In Fig. 1C and D, the central segments of the PRP profiles corresponding to the linker DNA involve a series of peaks of the same width and nearly equally spaced. This peaked structure is exclusively due to the DIM-2 dependent RIP because it completely disappears in the (rid+/+,dim-2Δ/Δ) background, but remains in the PRP profiles obtained in the (ridΔ/Δ,dim-2+/+) background [33] (see below). Similar peaks were also found in the PRP profiles of other constructs with this linker and varied repeat lengths [27], therefore, it should be determined by the linker base-pair sequence with little or no participation of the flanking DNA. In the DNA construct shown in Fig. 1A, as well as other previously used molecules of this series, the invariant segment comprising the linker and the left repeat copy was taken from the genome of *N. crassa*. Therefore, it might involve specific protein binding sites, and the periodical variations of the RIP propensity could be attributed to partial protection of DNA by some proteins not necessarily relevant to RIP.

To shed light upon this issue we compared two constructs that contained direct repeats of exactly the same length (802 bp) separated by exactly the same amount of linker DNA (729 bp). Their base-pair compositions are shown in the lower part of Fig. 4A. In the first construct (used earlier) the sequence had a uniform GC content of about 50%. In the second construct the repeated sequences were significantly enriched in AT-pairs. It was predicted that, because of a relatively weak homologous pairing, AT-rich repeats should be recognized less efficiently [29]. Therefore, a higher GC content was chosen for the linker sequence in order to increase the number of cytosine sites and amplify the RIP signal. It is known that, in vegetative cells of *N. crassa*, cytosines in AT-rich DNA are spontaneously methylated by DIM-2 via a pathway that also involves H3K9me3 methylation [37]. To check that the AT-rich sequence in our construct would not by a similar mechanism spontaneously induce DIM-2-mediated C-to-T mutations we also examined a supplementary construct that only carried one copy of this sequence rather than two (Appendix). No traces of RIP could be detected even in the rid+/+,dim-2+/+ background, thus confirming the notion that DNA homology is absolutely necessary for any RIP activity.

Fig. 4B compares the PRP profiles obtained for these two constructs subjected to DIM-2-mediated RIP in the absence of RID (ridΔ/Δ,dim-2+/+). It is seen that, despite different amplitudes due to the dissimilar DNA sequences in the linker regions, both profiles exhibit a similar series of peaks spaced apart by 115-120 bp. The PRP signals smoothly fall towards the borders, but some structure is also seen beyond the linker. Similar structures in the wild type profiles are overshadowed by the strong RID-dependent RIP activity. In Fig. 4C the central region is additionally compared with the wild type data for three different repeat lengths [27]. In these experiments the RIP intensity strongly varied (Appendix), but the fine PRP structure on the linker remained always similar, with some flattening of the third peak observed with shorter repeats.

**FIG. 4:**
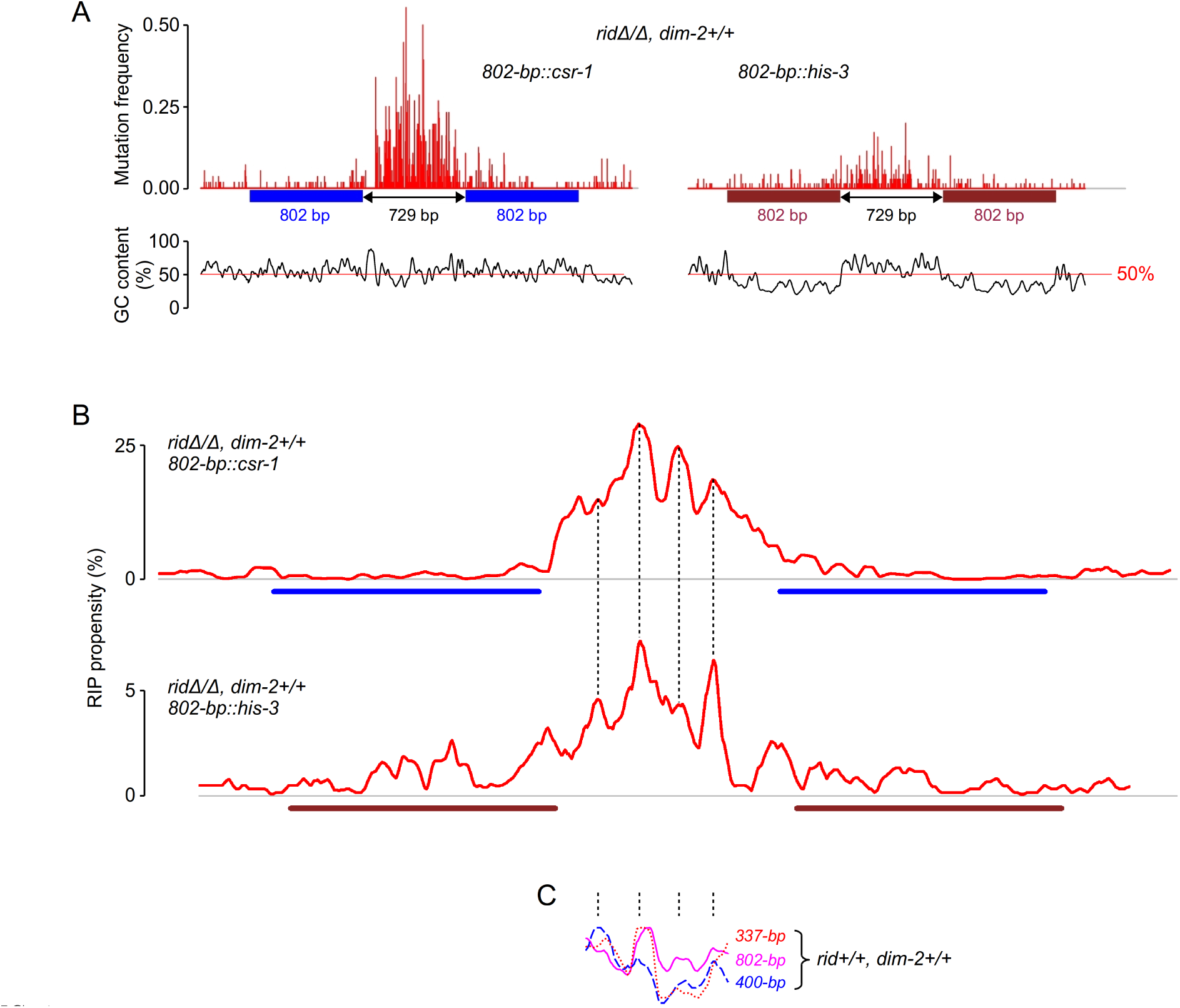
Characteristic fine structure of PRP profiles for the DIM-2 dependent RIP pathway. (A) Experimental mutation frequencies for two DNA constructs with contrasting GC-contents. The left construct was used previously [33]. The right construct with AT-rich repeats is described in Appendix. (B) Comparison of the PRP profiles. The vertical dashed lines mark the major similarly spaced peaks. The lower profile was shifted by about 50 bp. (C) Normalized PRP profiles obtained in the wild type *N. crassa* with the same linker and three different repeat lengths [27]. The plots were normalized on the corresponding maxima.

The DIM-2 dependent RIP requires the histone H3 lysine methyltransferase DIM-5 and the heterochromatin protein HP1 [33], which corresponds to the common mode of action of this DMT. All detected methylation in *N. crassa* depends on DIM-2 and DIM-5 [37], indicating that the activation of DIM-2 always occurs via its recruitment to the nucleosome. Therefore, the simplest reasonable interpretation of the regularly spaced peaks in Fig. 4B is that they result from DNA wrapping around some histone H3 complexes. A similar peak spacing observed for two DNA constructs with similar design and different sequences indicates that the linker length defines the total number of such structures whereas the base pair sequence defines their positioning on DNA. We will discuss this interpretation further below.

## DISCUSSION

### AT-rich repeats trigger RIP less efficiently

Our theoretical work [29] predicted that homologous AT-rich DNA segments should pair less effectively than those with higher GC content, suggesting that the AT-rich DNA would be a poor inducer of RIP. On the other hand, AT-rich sequences contain fewer cytosines that can potentially be mutated to thymines, i.e., these are also poor natural substrates of RIP. In the canonical RID-mediated RIP, DNA repeats act as both the inducers and the substrates for RIP, which complicates the testing of the above prediction. In contrast, in the DIM-2-mediated RIP, specifically in the context of a pair of closely-positioned direct repeats, the majority of mutations occur in the single-copy linker region between the repeats [33]. As a result, the two processes, namely, homology recognition and mutation, become decoupled. This feature allows examination of the ability of AT-rich repeats to be recognized as homologous, provided that the linker is sufficiently GC-rich to assure the RIP response.

A comparison of PRP profiles for the DIM-2-mediated RIP induced by repeats with strongly different GC content was shown in Fig. 4B. These results suggest that the AT-rich repeats indeed are much less efficient in inducing the DIM-2-dependent RIP. Alternatively, one might assume that the nearly five times lower RIP propensity in the second construct was due to an increased GC-content or just a different linker sequence. Such strong sequence effects can never be excluded, however, all our previous experiments revealed no indication that they can exist. Sequence effects on RIP are noticeable for partial homologies where the number of productive pairing modes is close to zero [27, 28], but they were not earlier observed for linkers and perfect repeats. Note also that in the second construct the RIP activity relatively easily spreads from the linker into the repeats, which is also attributable to weak binding of the paired homologs.

The observed difference between the RIP responses produced by the two types of sequences further supports the idea that the recognition of DNA homology for RIP occurs through direct homologous dsDNA-dsDNA binding during which GC pairs provide the principal contribution to the stabilization energy of the corresponding complexes [29]. According to this model, the formation of the both quartet types (involving pairing of AT or GC base-pairs) includes the creation of two additional hydrogen bonds, but the stabilization energy of the GC/GC quartets is much larger due to an unusually high electron polarization [29].

### DNA twisting and supercoiling

The mutation patterns in Fig. 4 show that the DIM-2-mediated RIP preferentially targets the single-copy DNA region between closely-positioned repeats. One can reasonably assume that, in this case, the DIM-2 pathway is specifically activated by the DNA/chromatin structure formed on the loop between the two paired DNA duplexes. The DIM-2-mediated RIP is particularly significant in the linker region between the direct repeats and it strongly depends upon the linker length (Fig. 3). It is also characterized by the regularly peaked structures (Fig. 4), with a maximum in the middle. All these features are virtually absent in the linker region between the inverted repeats, with the corresponding PRP profiles attributable to residual RID-dependent RIP near the repeat borders.

Some hints about the origin of these effects can be obtained from the analysis of the DNA topology in putative recognition complexes shown in Fig. 5. Consider the sketch of direct repeats in Fig. 5A. We suggested that the peaks in the PRP profiles analyzed in Fig. 4B represent a signature of some chromatin structures with DNA wrapped around histone cores. This is shown schematically by spheres. Taking into account the linker length and the number of resolved peaks in the PRP profiles one has to assume that the paired repeats are significantly bent. This bending may be assisted by other proteins omitted in this figure. According to our previous data the helical twist in paired repeats differs from that in free DNA [27]. In this case, the supercoiled configuration of the linker has to change because the helical linking number is constrained in the DNA loop closed by the two paired homologs. The homologous alignment is assumed to start from the region marked by the red oval. This first contact closes the loop that includes the linker and the two flanking segments of the repeats. The length of this closure is that of one repeat regardless of the site of the initial contact. The subsequent untwisting or over-twisting of the repeats must be compensated by supercoils on the linker.

**FIG. 5:**
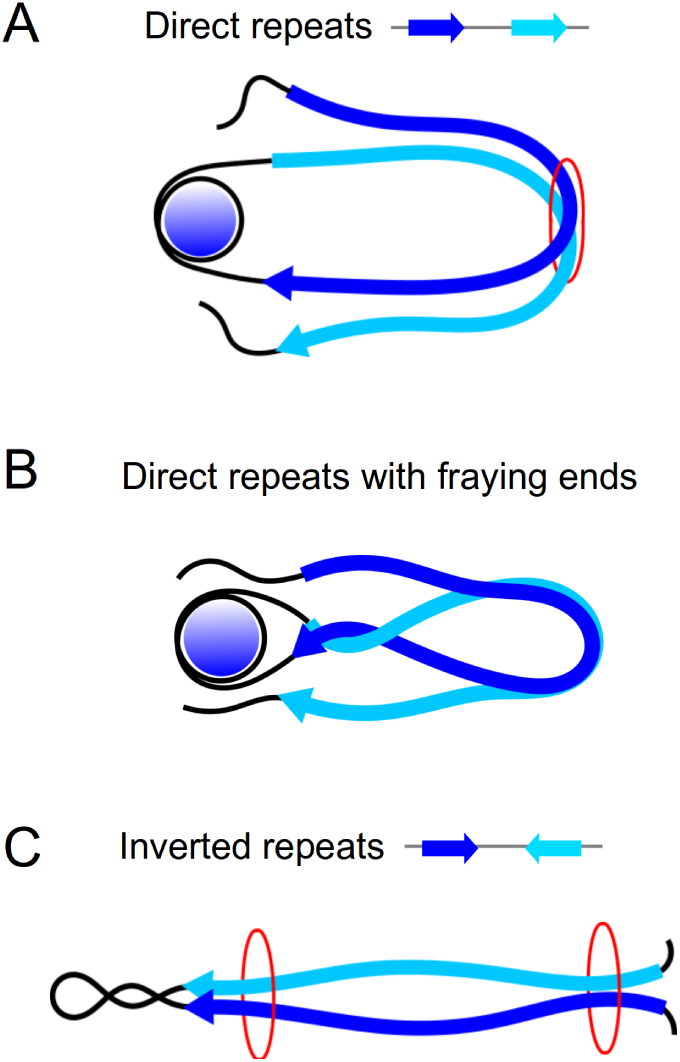
(A) The DNA topology in a putative homology recognition complex for a construct with direct repeats separated by a short linker. The repeat copies are shown by blue and cyan stripes with sequence alignments marked by arrows. The rest of the DNA is represented by thick black lines. Histone cores are shown by blue spheres. The early phases of pairing are shown, with red ovals marking alternative locations of sites of initial homology recognition. (B) A sketch of aligned direct homologs with fraying ends in a RIP-competent complex. (C) The DNA topology in putative recognition complexes formed by a construct with inverted repeats.

The number of helical turns exchanged between the linker and repeats can be significant. The most efficient RIP is observed with interspersed homologies corresponding to helical pitches of 11 and 12 base pair steps [27], which corresponds to a noticeable untwisting from the equilibrium B-DNA pitch of 10.5. At least 1.5 positive supercoils have to be added on the linker of the second construct in Fig. 3 to compensate for such changes in 330 bp repeats. One can predict that these effects should grow with the repeat length and fall with that of the linker. The first of these trends explains the variations in the third peak in the wild type profiles shown in Fig. 4C. The second trend accounts for the differences in the PRP profiles in Fig. 3 as well as the absence of similar DIM-2 dependent RIP on repeats separated by large genetic distances.

All earlier results concerning the mode of action of the DMT DIM-2 suggest that its activity depends on the presence of histone H3, the H3-lysine methyltransferase DIM-5, and the heterochromatin protein HP1 [33, 37]. This does not mean, however, that the linker DNA in the direct repeat construct in Fig. 5A must be negatively supercoiled and wrapped on octamer histone cores in standard nucleosomes. In fact, the members of the H3 histone family can participate in cores which differ by size, the length of DNA that they wrap and even the direction of wrapping [43]. At present, however, one cannot be sure about the sings and the amplitudes of the superhelical effects related with the homologous dsDNA-dsDNA paring because the initial untwisting of DNA can be further increased or inverted in the course of the subsequent binding of proteins involved in RIP.

The model displayed in Fig. 5A also sheds light on the intriguing systematic divergence of PRP profiles for paired repeats, which was shown in Fig. 2B. The shape of the PRP profile should depend upon the sequence as well as the particular structural environment of a given DNA segment. In the complex in Fig. 5A the homologous ends of repeats are found in different environments, namely, one is flanking the short stressed linker and the other is continued by “bulk” relaxed DNA. According to Fig. 2B this affects the RIP propensity in the boundary segments of about 150 bp. This DNA length is close to the threshold below which repetitive sequences are no longer recognized in RIP. This is also the length of the boundary zones of long repeats where RIP gradually disappears. One can reasonably suggest that all of these cases are related with weaker dsDNA pairing near the repeat ends, which should cause end fraying of aligned homologs. In the constructs considered here the fraying ends can be involved in the RID-dependent RIP by the mechanism sketched in Fig. 5B. When the homologous paring is partially broken, the supercoiling stress on the linker is redistributed to the flanking zones and results in formation of a plectoneme. The double helices in this structure can be erroneously recognized as paired homologs by proteins responsible for activation of the RID-dependent pathway. A certain number such proteins should be recruited to this region and we do not know yet which particular features of paired dsDNA are used for recognition. Two close and nearly parallel helices resemble paired homologs because the quadruplex recognition units are spaced by gaps of a few helical turns [29]. This similarity might be further increased by occasional matching of triplets in non-homologous helices.

A similar sketch of the recognition complex with inverted repeats is shown in the left panel of Fig. 5C. In this case the length of the similar loop depends upon the site of the initial homologous contact. If it occurs close to the linker its supercoiling is not affected during the subsequent stages. This situation is energetically favorable, therefore, even though the initial contact might occur at the opposite repeat end, a competition between these two pathways might favor the first one. Moreover, with this topology the linking number can be more readily maintained by forming plectonemes. Given this qualitative distinction from the structure in the right panel, one can surmise that the strikingly different RIP patterns of direct and inverted repeats in Fig. 3 can result from twisting of repeats during homologous pairing and formation of RIP-competent DNA-protein complexes.

The end faying can also occur in the context of inverted repeats. From the above considerations it follows that it should be smaller near the linker, and the RIP propensity is expected to be higher than at the opposite ends of the repeats. This agrees with the overall shape of the blue profile in Fig. 3. One can argue that, in this case, a sequence dependent decrease of RIP, if present, will have an approximate mirror symmetry and can produce the same effect. The last interpretation, however, disagrees with the data shown in the right panel of Fig. 3. These two profiles are taken from the flipped region marked in the middle panel, with the blue profile inverted. The inversion makes the base-pair sequences aligned, which results in matching positions of the main peaks evidently determined by the base-pair sequence. In contrast, the RIP propensity in both cases decreases with the distance from the linker, which agrees with the above structural interpretation.

### Possible structural role of repeat-induced DNA supercoiling

The proposed mechanism of repeat-induced supercoiling should be generally applicable beyond RIP. The DIM-2 dependent RIP pathway employs a series of proteins involved in the formation of constitutive heterochromatin in *N. crassa*. This finding led to the suggestion that RIP may represent a specialized case of the general mechanism involved in the initiation of heterochromatin on repetitive DNA in eukaryotic cells [33]. It is tempting to speculate that DNA supercoiling generated during the dsDNA-dsDNA pairing plays a conserved role and represents a structural signal used to program the behavior of chromatin. For example, long tandem repeats, such as those found in (peri-)centromeric regions of mammalian chromosomes, present numerous possibilities for creating looped DNA complexes like those in Fig. 5A. Assuming that dsDNA-dsDNA pairing requires partial untwisting of the two double helices [27], the positive superhelical density is expected to accumulate in the region between the paired dsDNA segments. This process should lead to the eviction of the canonical nucleosomes (which normally carry negatively supercoiled DNA) and can be facilitated by the association of proteins that favor positively supercoiled DNA, such as CenH3-containing hemisomes [44]. Once the paired repeats separate, their negative supercoiling can be relaxed by a topoisomerase IB, before the canonical nucleosomes are assembled back by a replication-independent mechanism [45], thus leading to the positive supercoiling density over the overall region involving the dissociated repeats and the intervening DNA (which in the case of a long array of tandem repeats also consists of repetitive DNA). This would explain the recently observed phenomenon of positive supercoiling of centromeric DNA repeats that have been subjected to replication in the reconstituted system in vitro [46]. Other scenarios can also be imagined. For example, in the absence of proteins that stabilize positive DNA supercoiling, the positive superhelical density in the linker region may be relaxed by topoisomerase IB, thus producing a region with negative superhelical density, once the paired repeats are dissociated. Similar mechanisms may be applicable in many situations and this hypothesis deserves further studies.

## AUTHOR CONTRIBUTIONS

A.K.M. designed and performed the theoretical part. E.G. carried out experiments. Both authors analyzed the data and wrote the article.

## ACKNOWLEDGEMENT

The work was supported by CNRS, the “Initiative d’Excellence” program from the French State (Grant “DYNAMO”, ANR-11-LABX-0011-01)”, grant GM044794 from the National Institutes of Health to Nancy Kleckner and research fellowships from The Helen Hay Whitney Foundation, The Howard Hughes Medical Institute, and the Charles A. King Trust to E.G.

## Appendix A Experimental Materials and Methods

### 1. Plasmid construction

Plasmid pEAG238A: the AT-rich fragment of the *AvHsp82* gene (corresponding to the nucleotide positions 33911-34710 of the GenBank entry EU637017.1) was amplified with the primers P238A-_F1 and P238A_R1 and inserted between the restriction sites *NotI* and *PacI* of the plasmid pMF272 targeted to the *his-3* locus [47].

Plasmid pEAG238B: a fragment of the plasmid pEAG238A containing the *AvHsp82* gene was amplified with the primers P238B_F1 and P238B_R1 and inserted between the restriction sites *EcoRI* and *ApaI* of the plasmid pEAG238A. The resultant construct contains two 802-bp repeats separated by a single-copy region of 729 base-pairs.

Annotated maps of pEAG238A and pEAG238B are provided as Supplementary Data (in GenBank format, individual plain-text files).

### 2. Manipulation of Neurospora strains

Linearized plasmid pEAG238A was transformed into the *rid+;his-3* strain FGSC#9720 [48]. Linearized plasmid pEAG238B was transformed into the *ridΔ;his-3* strain C02.1 [33]. Corresponding homokaryotic *his-3+* transformants (T503.4h carrying one copy of the *AvHsp82* fragment in the *rid+* background and T465.4h carrying two copies of the *AvHsp82* fragment in the *ridΔ* background) were purified by macroconidiation of the primary prototrophic transformants. The inserts were verified by PCR and sequencing. To examine the ability of the AT-rich *AvHsp82* fragment (when present in only one copy) to trigger any RIP mutation, transformant T503.4h was crossed (as a male parent) to the *rid+;his-3* strain FGSC#9539 [48]. To examine the ability of the duplicated AT-rich *AvHsp82* fragment to trigger RIP by the heterochromatin-related pathway, transformant T465.4h was crossed (as a male parents) to the *ridΔ;his-3* strain C03.1. All crosses were setup and analyzed exactly as described previously [33]. Spores were germinated on the minimal medium lacking histidine, thus selecting for prototrophic progeny corresponding to the “male” *his-3+* loci with the assayed constructs. 60 randomly selected “late” spores were analyzed for each condition.

### 3. Molecular analysis

Extraction of genomic DNA from germinated spores as well as PCR amplification, sequencing and assembly of the assayed constructs were done exactly as described previously [33]. The following primers were used for sequencing: NcHis3_F4.2, NcHis3_R1.2, GFP Seq5, GFP Seq6.

### 4. Primers used in this study

~~~
P238A_F1 AATAATGCGGCCGCCGATCAGGTACACGTGCATTTA
P238A_R1 CCTCCATTAATTAAGGCATTAATTCTTCGCAGTTTTCC
P238B_F1 AATAATGAATTCCAGGTACACGTGCATTTATGG
P238B_R1 AATAATGGGCCCAATTAAGGCATTAATTCTTCGCAG
NcHis3_F4.2 GTGGATGTCACAATGTCCCT
NcHis3_R1.2 CCCTAGATGATGGTTAGGCA
GFP_Seq5 CTTCAAGGAGGACGGCAACAT
GFP_Seq6 ACGCTGCCGTCCTCGATGTT
~~~

## Appendix B Computation of partitioned RIP propensity profiles

The propensity profiles discussed in the main text are meaningful under the following conditions: (i) the sequence dependence of RIP can be accounted for, (ii) the aggregate RIP intensity grows with the DNA length smoothly, and (iii) the accuracy and the sampling statistics are sufficient to observe along DNA the correlations between the density of RIP mutations and the corresponding density of cytosine sites. These three issues are considered below.

Fig. A1 demonstrates that, at least for CpA steps in *N. crassa*, the sequence context beyond the nearest cytosine neighbor has a minor effect. This was checked by using our previous data for direct repeats of different lengths [27]. To this end, the RIP probability *P* = *M*/*N* was measured where C is the number of mutations per cross and *N* is the number of sites. The primary RIP involves only 5-cytosine methylation and deamination to thymine, but the subsequent cycles of DNA replication produce daughter strands with G→A transitions. The total counts of C→T and G→A transitions, therefore, are summed up together and analyzed separately for cytosines with different neighbors. to compute partitioned probabilities *P*_CN_ where N denotes the other base in CpN and NpC steps, respectively.

**FIG. A1:**
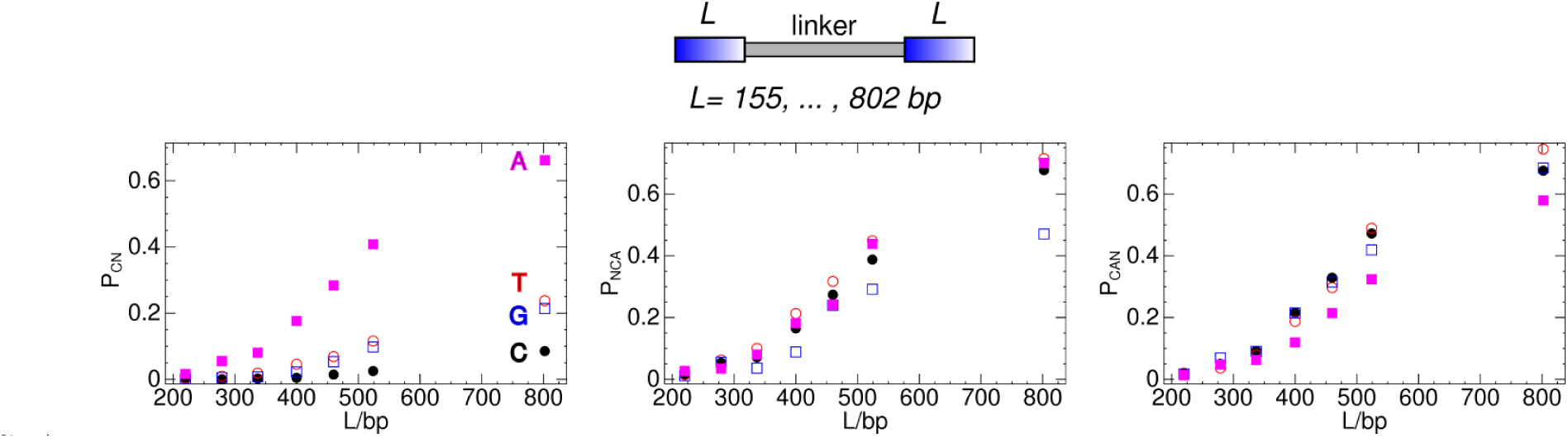
DNA length dependences of RIP probabilities for different types of cytosine duplets and triplets in the repeats. The identity of base N is indicated in the left panel. Symbols and colors are similar in all plots. The experimental data were taken from the earlier study [27]. The structure of the corresponding DNA constructs is shown on the top, with the blue gradient used for indicating the sequence homology and the mutual orientation of the repeats. The repeats are separated by a random DNA linker of 0.7 kbp.

There are eight cytosine duplets and eight triplets for every duplet. It is seen in Fig. A1 that the CpA steps indeed stand out, but the identity of the third nucleotide in triplets NCA and CAN is relatively unimportant. It is reasonable, therefore, to partition all cytosines into CpA, CpT, CpG, and CpC steps and assume that in the first approximation the intrinsic mutation rates do not depend upon other neighbors.

In addition, this figure illustrates a practically important feature that we call RIP saturation. With large repeat lengths, the RIP probabilities naturally converge to unity, which reduces all observable variations, including the sequence dependence. The maximal difference between CpA and other cytosine duplets reaches an order of magnitude under low RIP intensity.

The data shown in Fig. A1 also demonstrate that the partitioned RIP probabilities grow with the DNA length smoothly, which suggests that RIP can be characterized by the local densities of sites and mutations. Such densities, D(s) and C(s), can, in principle, be computed by moving a sliding window of an appropriate small length along DNA. However, this straightforward method results in rather low resolution because the analyzed sequences commonly contain regions with low site content. For the experimental data considered in the present study the minimal fixed window length is, in fact, around 80 bases, which is unnecessarily long. In our procedures we scan DNA with sliding windows of variable lengths so that the number of sites is fixed (usually 10). The site density at position s is that from the window with the closest midpoint. The short-range noise in the resulting profile is additionally reduced by smoothing with a sliding window of 20 bp. With this method, the resolution is proportional to the site density, while the latter is computed with a constant statistical error throughout the sequence.

The above approach is meaningful, however, only if the sampling statistics is sufficient to observe a linear correlation between the values of M(s) and D(s) computed from experimental data. This is checked for CpA and CpT duplets in Fig. A2 using the RIP frequencies for the 802 bp direct repeat construct from the earlier study (10). With good sampling statistics, the plots should exhibit linear trends, and the corresponding slopes should grow with the RIP intensity. Under exhaustive RIP, these plots converge to the square diagonal because every site can be mutated only once. Simultaneously, the RIP propensities should exhibit a uniform growth of uncorrelated fluctuations and convergence to the horizontal straight line. These fluctuations are not entirely statistical because the RIP propensity in the DNA regions with similar site densities differ due to structural factors and they can be revealed by studying profiles Pcn(s). The experimental data in Fig. A2 correspond well to the above pattern. The apparent residual correlations in the plots are due to incidental features like the effect of boundaries. Near repeat borders the RIP level naturally goes to zero, which results in apparent correlations if the site densities happen to reach extremal values.

**FIG. A2:**
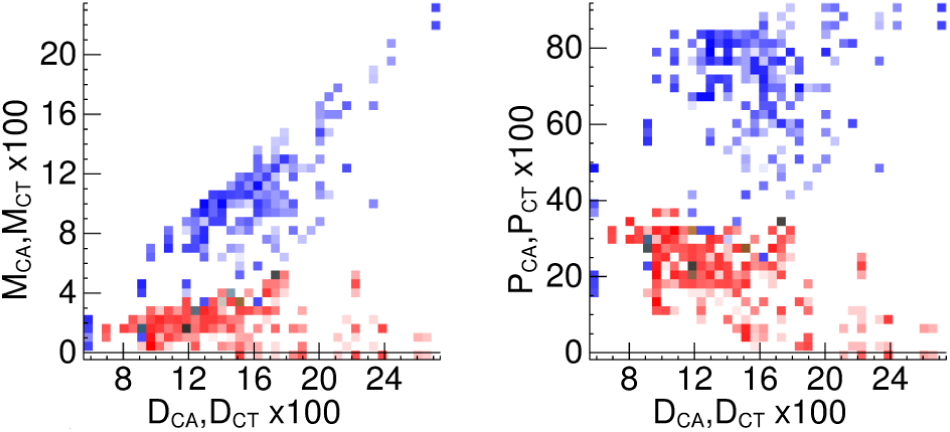
Correlations of the density of mutations and RIP propensities with the density of sites. The blue and red squares display correlations for ApA and ApT steps, respectively. Each square dot shows the number of regions of DNA with parameters indicated at the axes. This value is mapped to gradients of blue and red, with white corresponding to zero. The mutations were counted for both copies of the 802 bp repeats used in Fig. A1.

